# Investigating the importance of surface exposed loops in the gonococcal HpuB transporter for hemoglobin binding and utilization

**DOI:** 10.1101/2023.10.30.564842

**Authors:** Olivia A. Awate, Dixon Ng, Julie L. Stoudenmire, Trevor F. Moraes, Cynthia N. Cornelissen

## Abstract

*Neisseria gonorrhoeae* is the etiological agent of the sexually-transmitted infection gonorrhea and a global health challenge since no protective immunity results from infection and far fewer treatment options are available with increasing antimicrobial resistance. With no efficacious vaccines, researchers are exploring new targets for vaccine development and innovative therapeutics. The outer membrane TonB-dependent transporters (TdTs) produced by *N. gonorrhoeae* are considered promising antigen targets as they are highly conserved and play crucial roles in overcoming nutritional immunity. One of these TdTs, the hemoglobin transport system comprised of HpuA and HpuB, allows *N. gonorrhoeae* to acquire iron from hemoglobin (hHb). In the current study, mutations in the *hpuB* gene were generated to better understand the structure-function relationships in HpuB. This study is one of the first to demonstrate that *N. gonorrhoeae* can bind to and utilize hemoglobin produced by animals other than humans. This study also determined that when HpuA is absent, mutations targeting extracellular loop 7 of HpuB led to defective hHb binding and utilization. However, when the lipoprotein HpuA is present, these loop 7 mutants recovered their ability to bind hHB, although their growth phenotype remained significantly impaired. Interestingly, loop 7 contains putative heme binding motifs and a hypothetical α-helical region. Taken together, these results highlight the importance of loop 7 in the functionality of HpuB in binding hHb, and extracting and internalizing iron.

## INTRODUCTION

Gonorrhea, the common sexually transmitted infection (STI) that poses a serious global health issue, is caused by the etiological agent *Neisseria gonorrhoeae*, an obligate human pathogen. In 2020, the World Health Organization (WHO) estimated that 82.4 million new gonococcal infections occurred worldwide (1). In 2021, the Centers for Disease Control and Prevention (CDC) noted that 710,151 cases of gonorrhea were reported in the United States with a rate increase of 118% since 2009 (2). In 2018, the direct medical costs associated with gonococcal infections in the United States was about $271 million (3). A gonococcal infection often presents as urethritis or epididymitis in men or as cervicitis in women (4, 5). Approximately 80% of the cases in women are asymptomatic; an untreated gonococcal infection can lead to severe secondary sequelae such as infertility, ectopic pregnancy, pelvic inflammatory disease. Systemic infections in both sexes can lead to endocarditis, dissemination, and meningitis (5-9). One challenge with gonorrhea is that protective immunity is not conferred by a prior infection (4, 10, 11). Also, there has been a steady rise in gonococcal antimicrobial resistance (AMR) worldwide over the past 50 years (12), making the infection increasingly difficult to treat. With resistance to all classes of antibiotic used so far, the last method of treatment recommended by the CDC was a dual therapy with ceftriaxone and azithromycin (13-15). However, in 2020, due to the continuing surge in resistance, azithromycin use was discontinued, leaving ceftriaxone as the only treatment option, even though the occurrence of ceftriaxone resistance is also increasing (16-19). For this reason, researchers are now revisiting previously used antibiotics as possible alternatives to avert this AMR crisis (20). Thus far, there is also no effective vaccine against gonorrhea (21), and together with the declining availability of effective treatment options, it is anticipated that gonorrhea could ultimately be untreatable. We are, therefore, in dire need of new therapeutic options and preventive strategies against *N. gonorrhoeae*.

With no effective vaccines available, identifying promising targets for the development of vaccine and innovative therapeutics is crucial. The outer membrane TonB-dependent transporters (TdTs) are putative vaccine antigen targets as they are highly conserved among all pathogenic *Neisseria* strains (22), and most are not subject to high-frequency antigenic variation (23). More importantly, the TdTs play a vital part in allowing *N. gonorrhoeae* to overcome nutritional immunity, the process by which essential metal nutrients are restricted by the human host to hinder the growth of invading bacteria (24-26).

The hemoglobin receptor system, comprised of HpuA and HpuB, allows *N. gonorrhoeae* to acquire iron from human hemoglobin (hHb) (27, 28). The HpuAB system is phase variable (29, 30) but is turned on during active gonococcal infection in women during the first half of their menstrual cycle (31). Both HpuA and HpuB are thought to be required for utilization of hemoglobin–iron, in contrast the transferrin binding protein (TbpAB) system only requires expression of the TdT TbpA for iron utilization (22, 30) but the presence of TbpB can increase the efficiency of this uptake process (32). Protection studies using nonbinding mutant forms of the lipid-modified proteins TbpB and factor H binding protein (fHbp) as vaccine antigens have shown enhanced immunogenicity (33-35). While a similar phenomenon has not yet been described for the TdTs, it is possible that a cocktail of non-binding mutants derived from the TdTs could result in enhanced protection as was reported for two lipoproteins. Since there seems to be a selective advantage to expressing HpuAB during menses, HpuB mutants unable to bind and grow on hHb could potentially be considered as part of a protective vaccine cocktail against *N. gonorrhoeae*. We hypothesized that deletion of HpuB extracellular loops will reduce or abrogate Hb binding and consequently cause defect in growth with Hb as a sole iron source.

This current study aims to elucidate the interaction between gonococcal HpuB and hHb and to contribute to a detailed structure-function analysis of the gonococcal HpuAB system. HpuB extracellular loop deletion mutants were generated and analyzed for their binding ability to hHb and growth on hHb as a sole iron source, to better understand the binding and heme internalization mechanisms. In this current study, we found that mutations targeting extracellular loop 7, which contains conserved, putative heme-binding motifs, demonstrated impaired binding and growth, highlighting the importance of this loop in HpuB functions. We also found that the lipoprotein, HpuA, can restore Hb binding and iron internalization that is lacking in the HpuB loop7 mutant.

## RESULTS

### Hypothetical membrane topology of HpuB as a foundation for mutagenesis

Geneious was used to create a sequence alignment between meningococcal TbpA (for which the crystal structure is known [PDB 3V89]), gonococcal TbpA (WP_003693614) and HpuB (WP_003687084). Next, using the HpuB sequence in NetSurfP 2.0 and information about structural features of TbpA, the beta strands of HpuB were predicted. Finally, TOPO2 was used to create a transmembrane protein 2D topology and PHYRE2 was used to predict the folded structure of HpuB. A representation of the hypothetical HpuB topology model with extracellular loops, helices and plug domain is shown in Fig. 1. Using this model, the mutations were created to interrogate the role of each of the following predicted loop regions D236-246 (loop 2), aD306-311 (loop 3), D366-370 (loop 4), D538-544 (predicted helix in loop 7), DH548 and D555-559 (predicted conserved motifs in loop 7). While comparing the sequence of the gonococcal HpuB to that of other bacterial heme uptake systems, we identified 3 conserved motifs (Supp. Fig. 1), a histidine located between a FRAP motif and a NXXL (NPEL in *N. gonorrhoeae* FA19) motif. Additional single point mutants were generated by deleting either the histidine or the NPEL motifs but not the FRAP motif as it is predicted to be part of the barrel (36).

**Figure 1:**
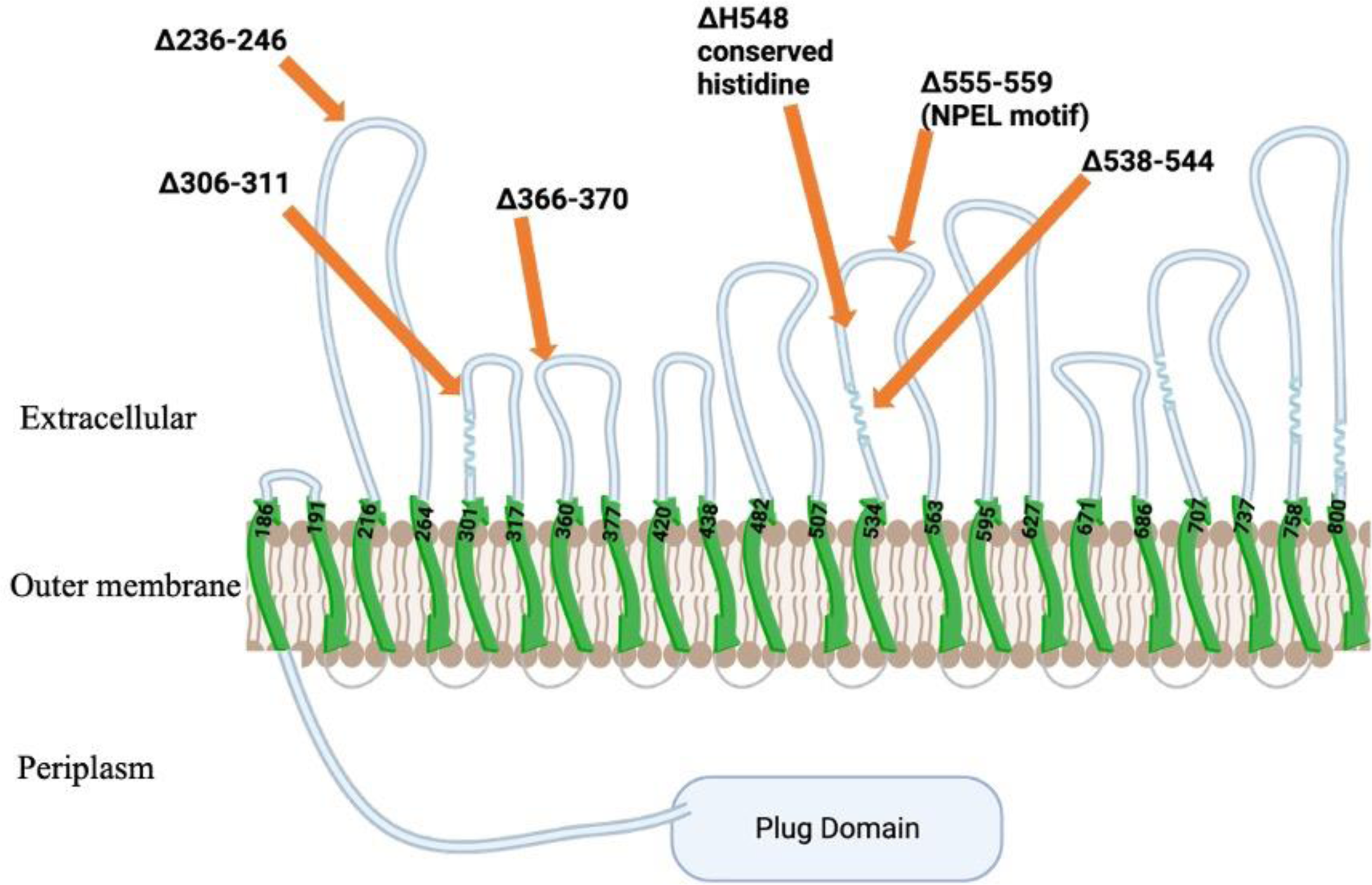
Hypothetical topology map of HpuB as a foundation for mutagenesis. The putative loops are extracellularly located with loop helices represented by coil-like regions. Beta strands are shown in green within the outer membrane. The orange arrows indicate the locations of the deletion mutations generated in this study. Image created with BioRender.com.

Mutated *hpuB* genes were created and confirmed to be expressed from an inducible promoter in an ectopic site in the gonococcal chromosome. The *hpuA* and *hpuB* genes are natively expressed only under iron-restricted growth conditions (30, 37). To avoid fluctuations in protein expression levels under iron restriction, we constructed two strains mutated in the native locus; one was unable to produce either HpuA or HpuB (RSC150). The other mutant strain had a locked, phase-on *hpuA* gene and an inactivated *hpuB* gene in the native chromosomal site (RSC275) (Table 1). Plasmid pVCU234, a complementation vector with an isopropyl β-d-1-thiogalactopyranoside (IPTG) inducible promoter followed by a strong ribosome binding site, was used to ectopically insert wild-type (WT) or mutated versions of *hpuB* into the RSC150 (A- B-) or RSC275 (A+B-) backgrounds (see Fig. 2A and B for schematic). Using this approach allowed for better control over the production of HpuB, with the addition of 1 mM IPTG, rather than iron restricted growth conditions.

**Figure 2:**
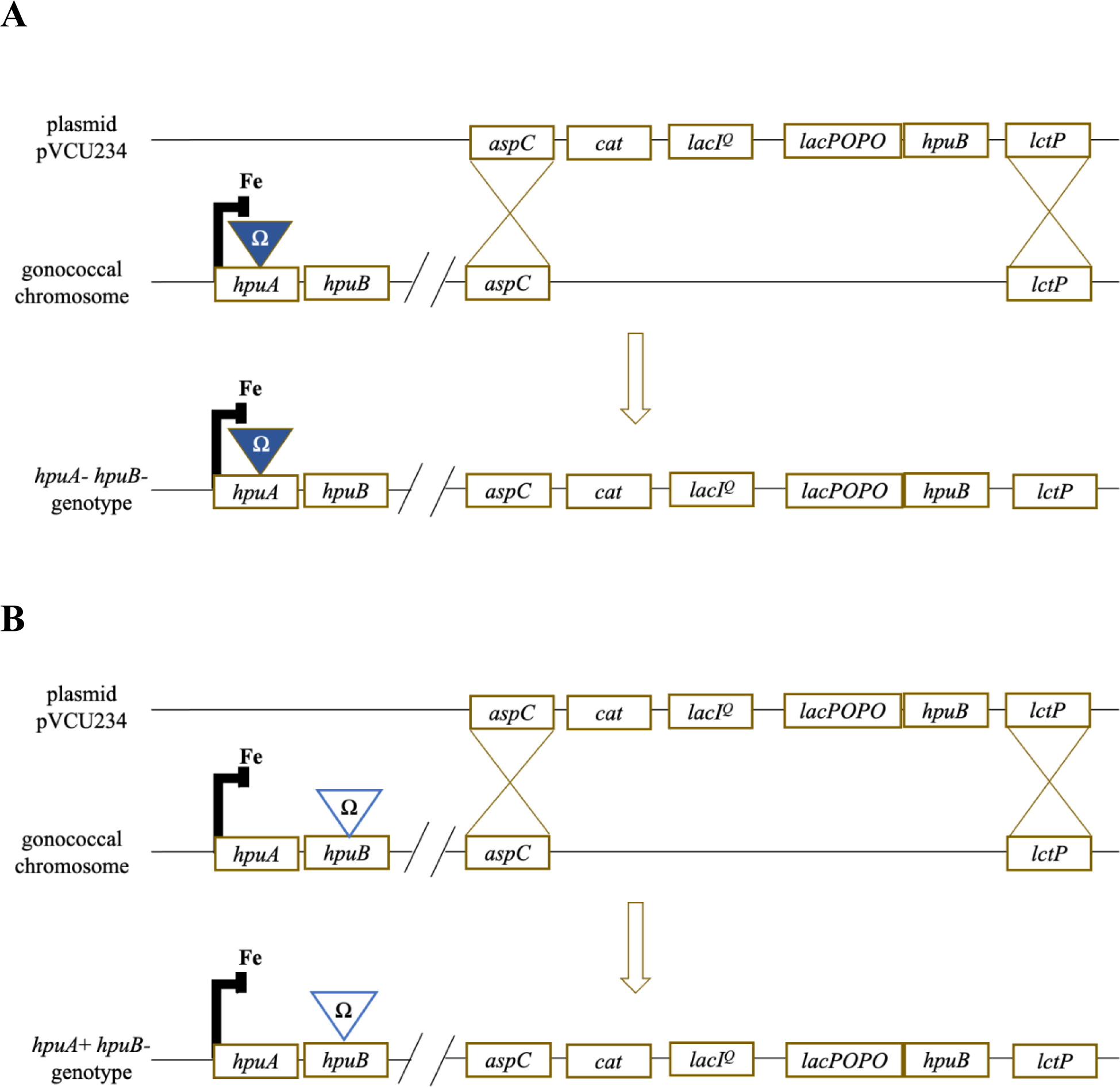
Diagram demonstrating how *hpuB* mutations were moved into the gonococcal chromosome generating gonococcal strains that were either HpuA-(panel A) or HpuA+ (panel B). Wild type and mutated versions of *hpuB* were cloned into a complementation vector, pVCU234, behind an IPTG inducible promoter and ribosome binding site. Transformation of *N. gonorrheae* results in wild-type or mutated versions of *hpuB* being inserted into an ectopic site between the *aspC* and *lctC* loci. The chromosome of the recipient strain is either *hpuA-hpuB-*, i.e RSC150 (panel A) or *hpuA+ hpuB-*, i.e RSC275 (panel B). *lacPOPO* represents the IPTG inducible promoter and Fe represents the iron repressed promoter.

**TABLE 1.** Bacterial Strains and Plasmids used in this Study.

| Strain or Plasmid | Description | Source or reference |
| --- | --- | --- |
| Strains |  |  |
| *E. coli* |  |  |
| TOP10 | *F2 mcrA D(mrr-hsdRMS-mcrBC) f 80lacZDM15 DlacX74 nupG recA1 araD139 D(ara-leu)7697 galE15 galK16 rpsL(Strr) endA1 l2* | Invitrogen |
| Gonococcal |  |  |
| RSC150 | FA19 *hpuA::Ω* (*hpuA*- *hpuB*-) | This Study |
| RSC275 | FA19 *hpuB::Ω* (*hpuA*+ phase one, *hpuB*-) | This Study |
| RSC276 | RSC150 + ectopic WT *hpuB*(*hpuA*- *hpuB*+) ) | This Study |
| RSC277 | RSC150 + ectopic Δ236-246 mutant *hpuB* (loop2) | This Study |
| RSC278 | RSC150 + ectopic Δ306-311 mutant *hpuB* (loop3) | This Study |
| RSC279 | RSC150 + ectopic Δ366-370 mutant *hpuB* (loop4) | This Study |
| RSC281 | RSC150 + ectopic Δ538-544 mutant *hpuB* (loop7 helix) | This Study |
| RSC282 | RSC150 + ectopic Δ555-559 mutant *hpuB* (loop7 NPEL motif) | This Study |
| RSC283 | RSC150 + ectopic ΔH548 mutant *hpuB* (loop7 Histidine motif) | This Study |
| RSC284 | RSC275 + ectopic WT *hpuB*(*hpuA*+ *hpuB*+) ) | This Study |
| RSC285 | RSC275 + ectopic Δ236-246 mutant *hpuB* (loop2) | This Study |
| RSC286 | RSC275 + ectopic Δ306-311 mutant *hpuB* (loop3) | This Study |
| RSC287 | RSC275 + ectopic Δ366-370 mutant *hpuB* (loop4) | This Study |
| RSC289 | RSC275 + ectopic Δ538-544 mutant *hpuB* (loop7 helix) | This Study |
| RSC290 | RSC275 + ectopic Δ555-559 mutant *hpuB* (loop7 NPEL motif) | This Study |
| RSC291 | RSC275 + ectopic ΔH548 mutant *hpuB* loop7 Histidine motif) | This Study |
| Plasmids |  |  |
| pVCU234 | pKH37 1 ribosome-binding site | (66) |
| pGSU442 | pVCU234 + WT *hpuB* | This Study |
| pGSU443 | pVCU234 + Δ236-246 mutant *hpuB* | This Study |
| pGSU444 | pVCU234+ Δ306-311 mutant *hpuB* | This Study |
| pGSU445 | pVCU234+ Δ366-370 mutant *hpuB* | This Study |
| pGSU447 | pVCU234+ Δ538-544 mutant *hpuB* | This Study |
| pGSU448 | pVCU234+ Δ555-559 (NPEL motif) mutant *hpuB* | This Study |
| pGSU449 | pVCU234+ ΔH548 (conserved histidine) mutant *hpuB* | This Study |

After growing the *hpuB* mutants with or without IPTG, a western blot was performed to confirm the production of HpuB in the presence of IPTG (Fig. 3). Strain RSC150 (A-B-) was used as a negative control whereas RSC284 (A+B+) with a native and locked phase-on *hpuA* gene and an ectopic *hpuB* gene, was used as a positive control. A stain free gel confirmed that the lanes were loaded consistently. Immunoblot analysis using a HpuB peptide-specific polyclonal antiserum (ordered from Biosynth), confirmed that HpuB production was regulated by IPTG addition in all the mutant strains. Following this confirmation, we progressed to characterization of the binding and growth phenotypes of these mutants.

**Figure 3:**
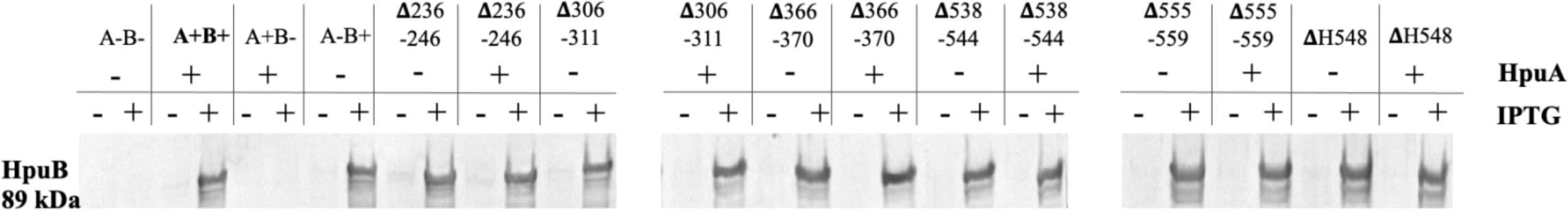
Mutated *hpuB* genes were created and confirmed to be expressed from an inducible promoter in an ectopic site in the gonococcal chromosome. HpuB mutant strains were grown on GCB plates containing DFO (Fe limiting), with (+) or without (-) 1 mM IPTG. Cells were collected from plates and resuspended into PBS. From these suspensions, whole-cell lysates were prepared by standardizing the cell suspensions to an OD_600_ of 1, pelleting cells and resuspending pellets in lysis buffer. Western blots were performed to characterize the production of HpuB in the presence or absence of IPTG using anti-HpuB antibody. The strain with WT *hpuB* gene in the ectopic site and native *hpuA* gene was used as a positive control (A+B+). The strain lacking both native *hpuA* and *hpuB* genes (A-B-) was used as a negative control. Equal loading of each lane is confirmed by examination of stain-free gels before electro-transfer for western blot (not shown). Data shown is representative of 3 biological replicates.

### Loop 7 *hpuB* mutants were impaired for growth on hemoglobin as a sole iron source

We evaluated whether any of the mutations impacted the ability of *N. gonorrhoeae* to utilize hemoglobin and whether the presence or absence of the lipoprotein, HpuA, also impacted utilization. We performed growth assays in metal-restricted conditions, with Desferal (DFO, an iron chelator) and IPTG to induce HpuB expression before adding 0 or 1 µM human hemoglobin (hHb) as the sole source of iron. The growth of all the mutant strains, under iron restriction and with the presence of IPTG, was compared to that of the positive control A+B+ (Fig. 4). In the absence of any iron source (0 µM hHb), no growth was observed for any of strains regardless of the expression of native *hpuA* (Supp. Fig. 2A and B). However, when 1 µM hHb was added, the *hpuB* mutants demonstrated different levels of growth. As expected, the negative control, RSC150 (A-B-), exhibited no detectable growth and the positive control, RSC284 (A+B+), demonstrated the most robust growth.

**Figure 4:**
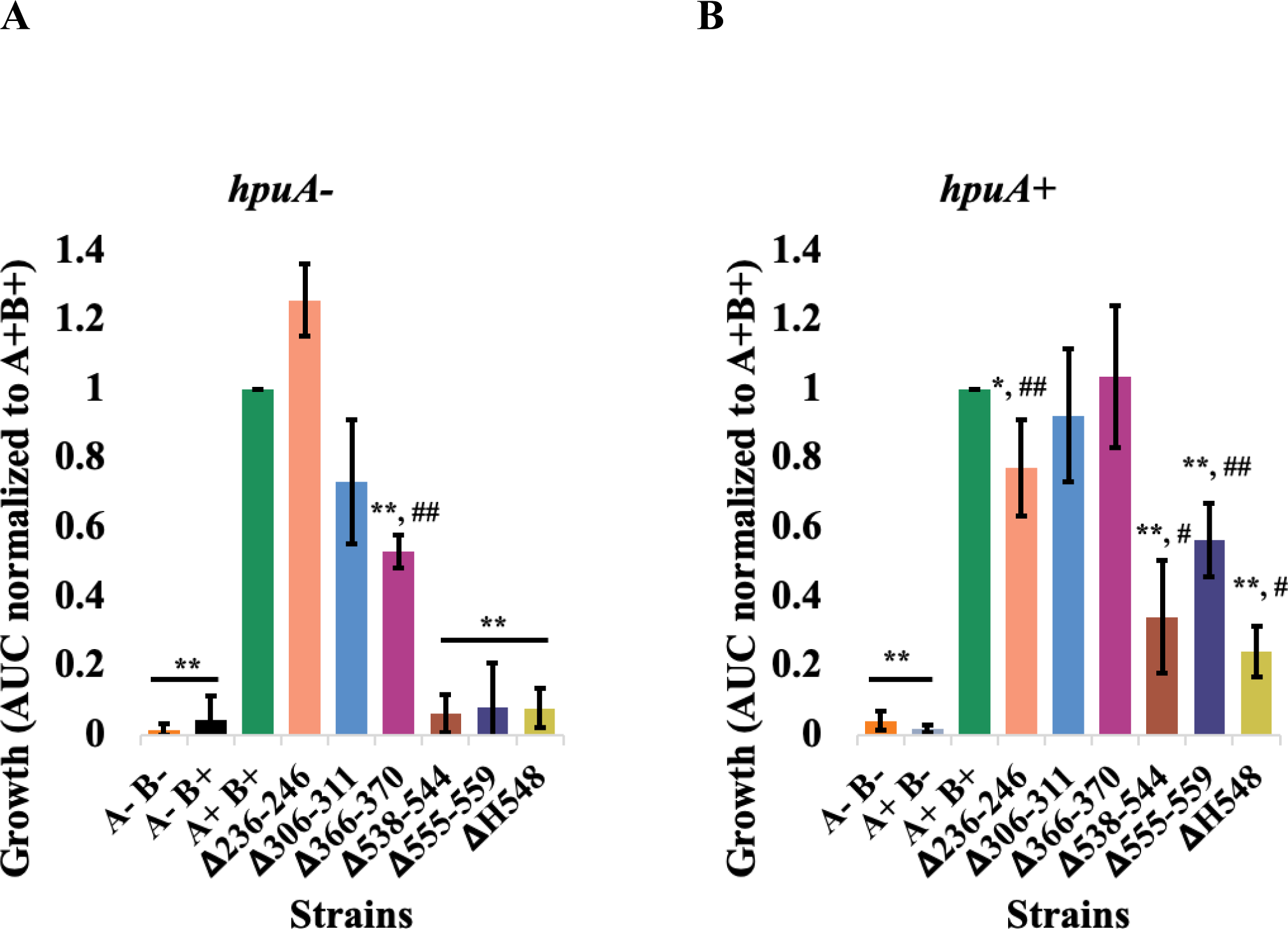
Loop 7 HpuB mutants are impaired for growth on hemoglobin as a sole iron source. Gonococci were grown on GCB/DFO/IPTG plates before being resuspended in CDM. The cell suspensions were standardized to an OD_600_ of 0.002 before being added to a 96-well plate containing DFO, IPTG and 1 µM hHb. Cells were grown for 21 hours, during which the OD_600_ was recorded in 30-minute intervals to assess the growth of strains in the mutant *hpuA* background (A) and in the wild-type *hpuA* background (B). A-B-is used as a negative control and A+B+ as a positive control. A+B-as well as A-B+ also represent controls as both proteins are thought to be required for growth. From the growth curves, the area under the curve (AUC) was calculated using GraphPad Prism then normalized to growth by the A+ B+ strain. The data from three biological replicates were analyzed to generate the means and standard deviations shown. Statistically significant differences were identified by using the Student’s *t-test* relative to the A+B+ strain (*, p < 0.05; **, p < 0.005) and to A-B-strain (#, p < 0.05; ##, p < 0.005).

When HpuA is not expressed, the D236-246 and D306-311 mutants that demonstrated growth levels similar to the A+B+ strain; D366-370 was both significantly lower than A+B+ and higher than A-B-. The loop 7 mutants D538-544, DH548, D555-559 all demonstrated a significant growth defect, comparable to the A-B-strain (Fig. 4A). HpuA and HpuB are both thought to be required for hHb utilization (30). To assess the requirement for HpuA in the mutant strains generated in this study, we measured the growth of RSC275 (A+B-) and RSC276 (A-B+), and both strains grew to the extent of the A-B-strain (Fig. 4A). This observation supports the importance of producing both HpuA and HpuB for hHb utilization.

In strains where HpuA was expressed, the D306-311 and D366-370 mutations resulted in growth levels similar to the A+B+ strain. In contrast the D236-246 strain grew significantly below that of the A+B+ strain but still higher than A-B-. The loop 7 mutants, D538-544, D555-559, DH548, demonstrated a significant decrease in growth although they recovered some growth compared to when HpuA is not expressed (Fig. 4B).

### Loop 7 *hpuB* mutants recover hHb binding ability when HpuA is produced

To determine the binding phenotypes of these *hpuB* mutants, we performed whole-cell ELISA. As shown in Fig. 5, various levels of hHb binding were observed for the mutant strains. As expected, the negative control, RSC150 (A-B-), showed the least detectable binding and the positive control, RSC284 (A+B+), showed the highest level of binding. When HpuA was absent, D236-246 and D306-311 mutants bound hHb to the level of the A+B+ strain, while D366-370 demonstrated significantly reduced hHb binding. The loop 7 mutants D538-544, D555-559, DH548 exhibited profound hHb binding defects, comparable to that detected with the A-B-strain (Fig. 5A). RSC276 (A-B+) also bound hHb at the level demonstrated by the A+B+ strain, suggesting that HpuB alone can accomplish binding of hHb in this system.

**Figure 5:**
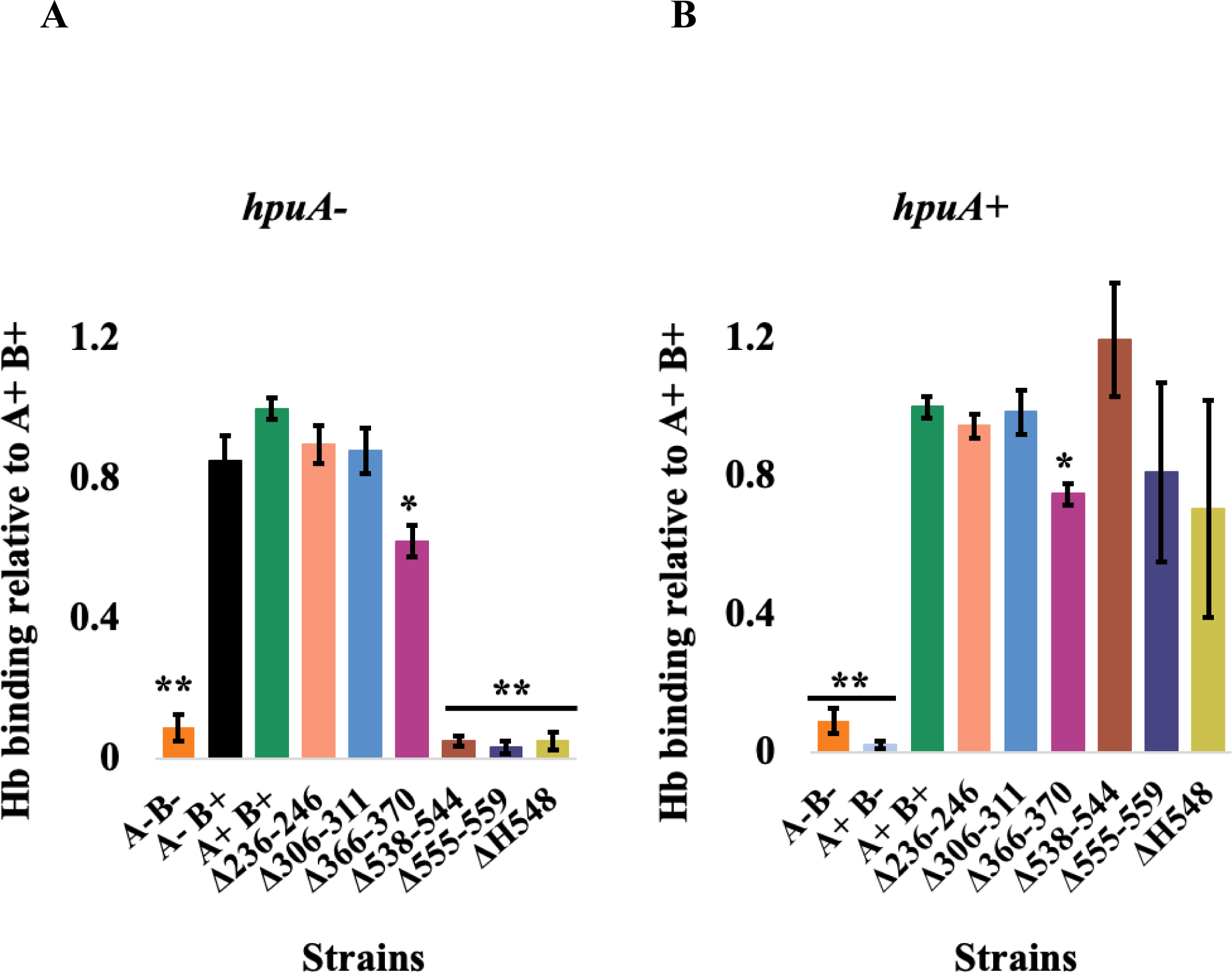
Loop 7 HpuB mutants recover hHb binding ability when HpuA is produced. Gonococcal HpuB mutants were grown on GCB/DFO/IPTG plates before being resuspended in PBS to an OD_600_ of 1. Cell suspensions were added to a 96-well ELISA plate and allowed to dry prior to blocking. The ELISA plate was then probed with HRP-conjugated hHb before being washed and developed with a TMB substrate. The binding ability of strains without (A) and with (B) native *hpuA* was assessed. The absorbance at 450 nm was read to quantify the signal. A-B-is used as a negative control and A+B+ as a positive control. A+B-as well and A-B+ are also used as controls since both HpuA and HpuB are thought to be required for binding. All the strains were normalized to A+B+ and showed as a percentage of A+B+. Three biological replicates are represented with their means and standard deviation shown. A Student’s *t-test* was used to assess the statistically significance differences relative to A+B+ (*, p < 0.005; **, p < 0.0005).

When the lipoprotein HpuA was expressed in the native site, all mutants, except for D366-370, demonstrated hHb binding levels similar to the A+B+ strain. The D366-370 mutation resulted in a strain demonstrating a significant decrease in binding when compared to the A+B+ strain (Fig. 5B). RSC275 (A+B-) showed no detectable hHb binding, indicating that production of HpuA alone does not allow hHb binding.

### Mutated HpuB variants are expressed on the gonococcal cell surface and susceptible to trypsin digestion

To ensure that the mutations generated did not affect the surface exposure and the overall conformational fold of the resulting HpuB proteins, we conducted a cell-surface protease digestion. The mutated strains were grown on plates supplemented with both DFO and IPTG. Following overnight growth, colonies were recovered from plates and resuspended in PBS; standardized cell suspensions were subjected to a time course of trypsin digestion (38) before whole-cell lysates were prepared. The lysates were then subjected to SDS-PAGE and transferred to nitrocellulose for subsequent detection of HpuB and proteolysis products using the HpuB peptide-specific polyclonal antiserum (Fig. 6). It should be noted that only the proteolytic products containing the peptide epitope, used to generate the antiserum (see methods and materials), were detected. A proper extracellular localization is confirmed by having a digestion pattern similar to the A+B+ wild-type strain. Over the time course utilized, the full length HpuB (∼89 kDa), was digested into ∼ 65kDa, ∼50 kDa, and ∼25 kDa fragments as seen in the A+B+ control strain. Mutations at locations D236-246, D306-311 and D366-370 resulted in HpuB-producing strains that display digestion patterns identical to that of the A+B+ control, indicating proper folding and surface exposure. The loop 7 mutants, however, demonstrated more proteolytic products than the A+B+ strain, with additional fragments at ∼55 kDa, ∼44 kDa, and ∼30 kDa, consistent with surface presentation but also additional exposure of trypsin sensitive digestion sites (Fig. 6). Taken together, these data indicate that all the mutants generated in this study presented HpuB on the gonococcal cell surface.

**Figure 6:**
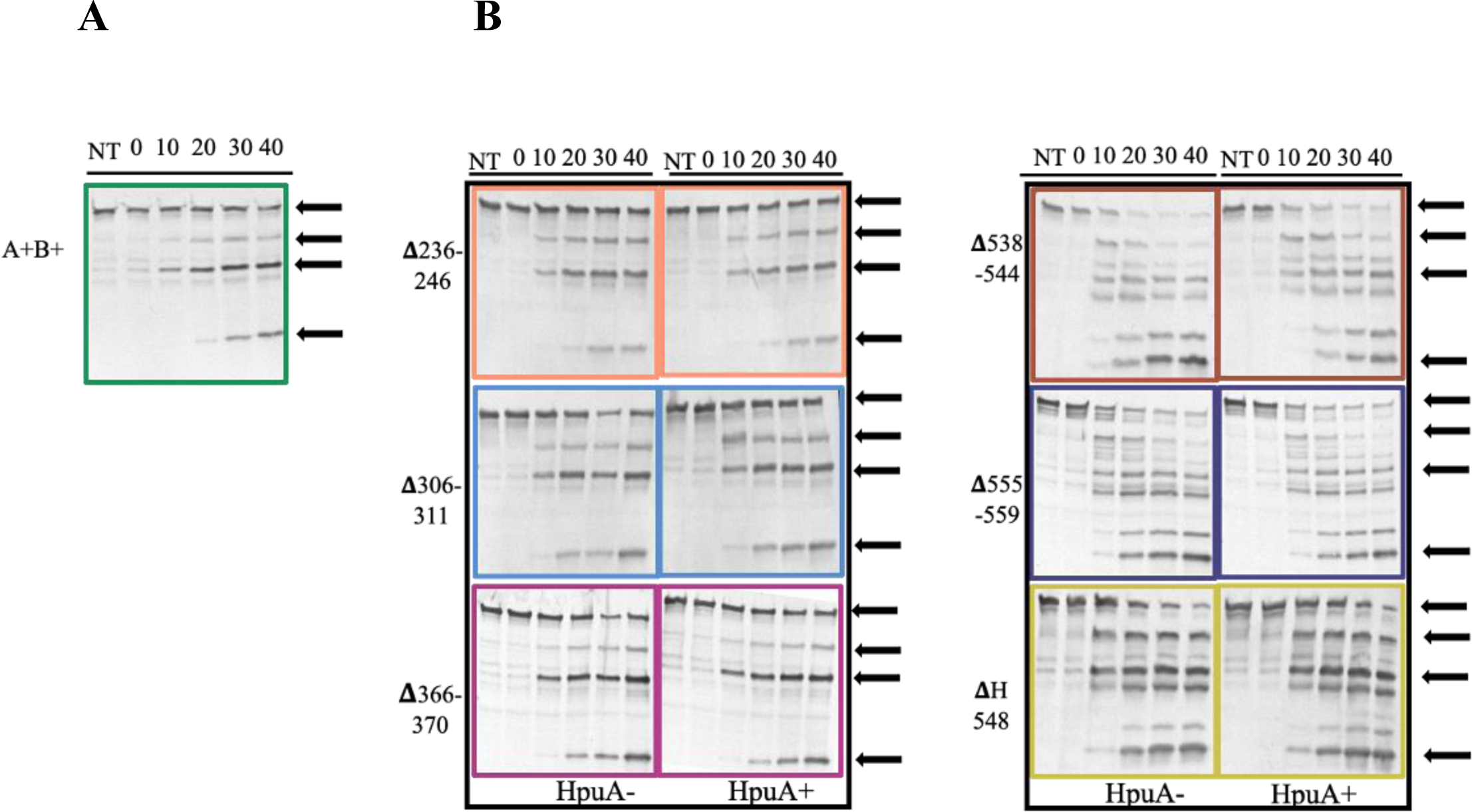
Mutated *hpuB* variants are expressed on the gonococcal cell surface and susceptible to trypsin digestion. Strains were grown on GCB/DFO/IPTG plates and resuspended in PBS to an OD_600_ of 0.4. Iron-starved whole cells were then treated with trypsin for 0, 10, 20, 30 and 40 min before the reaction was stopped with aprotinin. NT represents no treatment. Next, the cells were pelleted, and the lysates were subjected to SDS-PAGE and western blot. The blots were probed with anti-HpuB antibody followed by an AP-conjugated IgG secondary antibody. NBT/BCIP was used to develop the blots. A+B+ shows the positive control trypsin digestion pattern (A). The digestion pattern of the mutants is shown (B). The black arrows indicate the proteolytic products seen in the positive control.

### The deletions in loop 7 moves histidine 548 further away from the closest heme **group.**

After obtaining the results described above, we sought to better understand the role of loop 7 and in particular the role of the conserved histidine and the NPEL motif, which are conserved among heme transporters of many Gram-negative bacteria. To analyze the interfaces in a HpuA-HpuB-Hb ternary complex, the Alphafold2-based Colabfold running in Alphafold-multimer mode (39-41) was used to generate predictive models of the three proteins in complex. A known Hb-heme crystal structure (PDB:1hho) was super-imposed onto the complex to correctly place the heme groups since ligands cannot be generated in the modeling software. Our predicted ternary structure shows HpuB in green, HpuA in blue, Hb in red, and the corresponding heme group in white with its Fe^2+^ as a red ball (**Fig. 7A**). A zoomed-in image of the interaction shows HpuB in green with residue H548 highlighted in red and the distance between the closest heme group and H548 is measured in angstroms (Å); the distance between the closest heme group and H548 is 4.6 Å (**Fig. 7B**). However, this distance increases to 18.1 Å when the predicted loop helix D538-544 is deleted (**Fig. 7C**) and to 22.3 Å when the NPEL motif D555-559 is deleted (**Fig. 7D**). Interestingly, in the predicted binary complex of HpuB and Hb only, the distance between residue H548 and the heme group is 5.7 Å, in the mutants this distance is 14.2 Å when D538-544 is deleted and 10.5 Å when D555-559 is deleted (Supp. Fig. 3). The relative position of the histidine, thought to be essential for receptor function (42), changes with the D538-544 and D555-559 deletions, pushing the histidine further away from the surface and from interacting with the heme group in the hHb.

**Figure 7:**
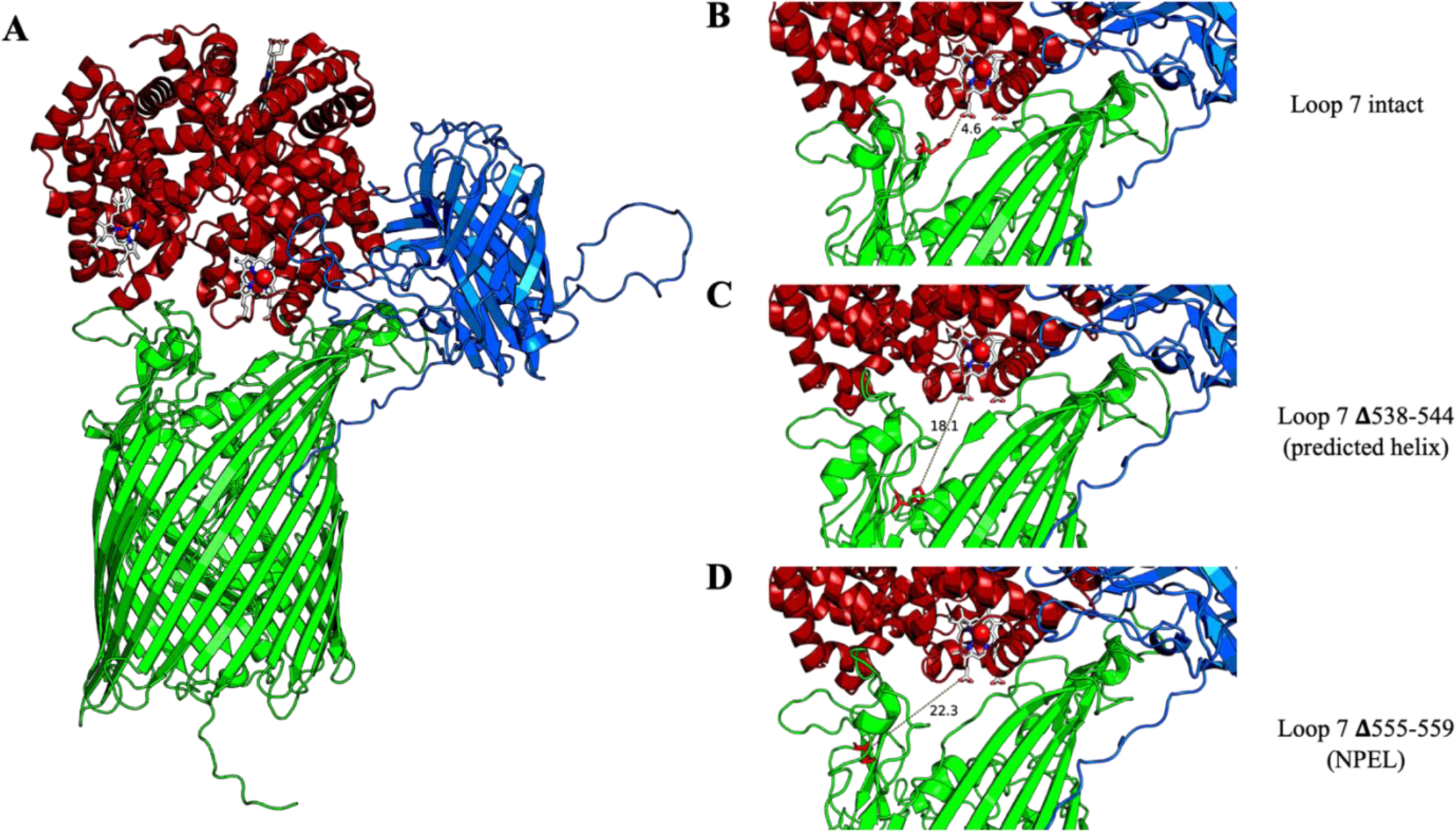
The deletions in loop 7 move histidine 548 further away from the closest heme group. HpuA-HpuB-Hb structure. AlphaFold2 was used to get a prediction model of the structures. A known Hb-heme crystal structure (PDB:1hho) was super imposed to show the heme groups. HpuB is in green with H548 highlighted in red, HpuA in blue, Hb in red, heme group in white with is Fe^2+^ as a red dot. The distance in angstroms (Å) between the heme group and histidine at position 548 is measured.

### *N. gonorrhoeae* can bind and utilize hemoglobin produced by animals other than humans, as a sole iron source

To date, *N. gonorrhoeae* has only been demonstrated to bind and utilize TdTs ligands primarily of human origin (43, 44) To determine if this concept applies to the HpuAB system and whether *N. gonorrhoeae* only binds and utilizes hemoglobin of human origin, we performed a competition assay and a growth assay. A wavelength scan (data not shown) was also conducted to ensure that the purchased Hb from all species of origin, were in the reduced (ferrous iron) form.

A competition assay was conducted to establish whether *N. gonorrhoeae* can bind to Hb from mouse (mHb), pig (pHb) and rat (rHb). RSC284 (A+B+) was grown as described above and tested by whole-cell ELISA for hHb binding in the presence of other species of Hb as competitors. As expected, the lowest detectable level of binding was seen in the negative control (blocker only) and the highest level of binding was detected when hHb-HRP was allowed to bind without competitor (Fig. 8). Unlabeled human, mouse, pig, and rat hemoglobin all competed with HRP-conjugated hHb, as seen by a significant decrease in labeled hHb binding. *N*. *gonorrhoeae* can, therefore, bind mouse, pig, and rat Hb in addition to hHb.

**Figure 8:**
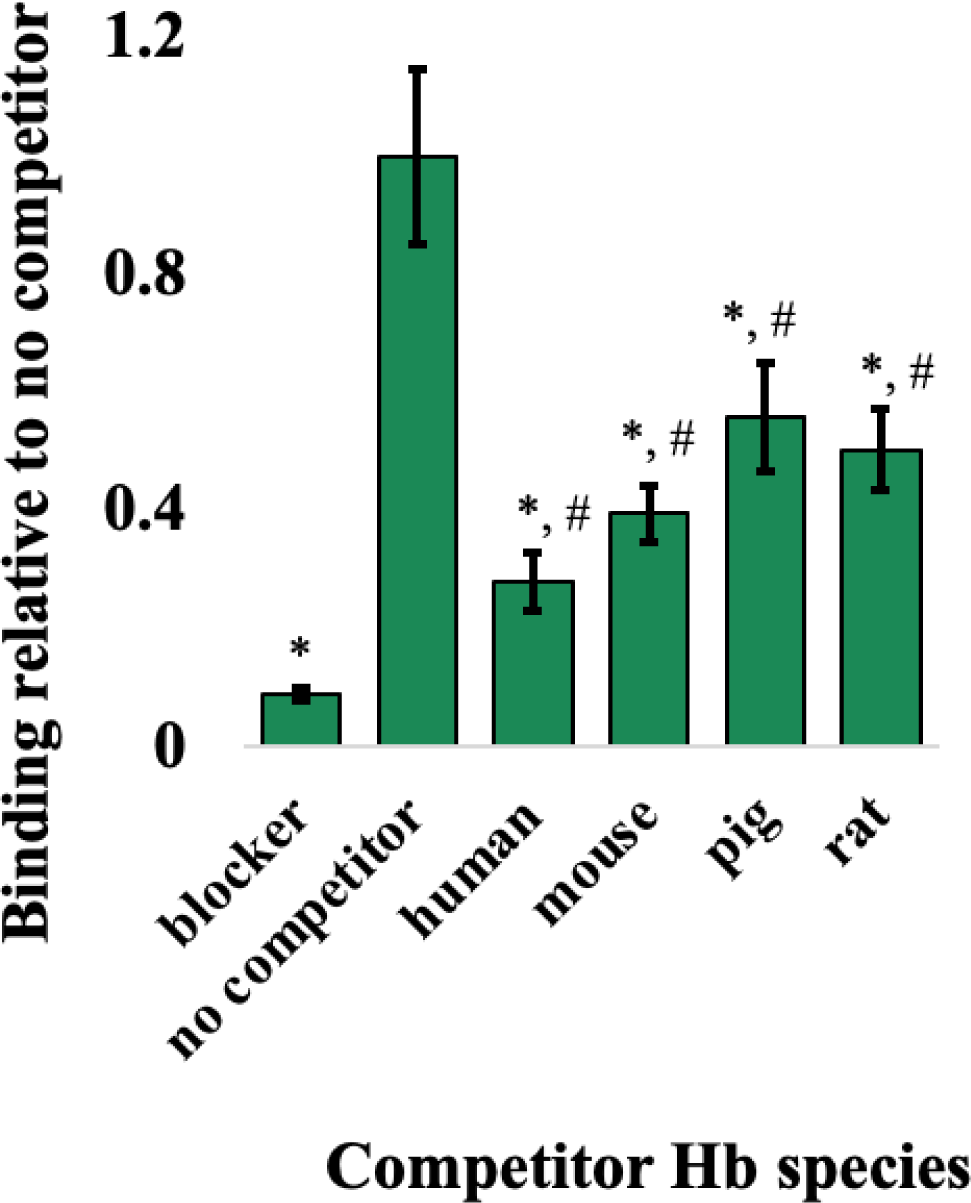
*N. gonorrhoeae* can bind hemoglobin produced by animals other than humans. A competition assay was used to establish whether *N. gonorrhoeae* can bind to Hb from mouse (mHb), pig (pHb) and rat (rHb). A+B+ was grown on GCB/DFO/IPTG plates before being resuspended in PBS to an OD_600_ of 1. The cell suspension was added to a 96-well ELISA plate and allowed to dry prior to blocking. The ELISA plate was then probed with either no Hb (i.e blocker), 5 nm hHb-HRP without competitor or 5 nm hHb-HRP + 20X (100 nm) excess competitor. The competitors used were unlabeled hHb, mHb, pHb or rHb. Next, step was to develop with TMB. 5 nm hHb-HRP + no competitor represents the positive control, and the blocker condition represents the negative control. Statistically significant differences were assessed by the Student’s *t-test* relative to the no competitor condition (*, p < 0.0005) and the blocker condition (#, p < 0.0005). The means and standard deviation of 4 biological replicates is shown.

Given this result, we next assessed the ability of the gonococcus to utilize Hb from different species by conducting a growth assay as described above, but this time, with either hHb, mHb, pHb or rHb provided as the sole source of iron. The growth of RSC284 (A+B+) with Hb from different species was compared to the growth of A+B+ in the presence of hHb (Fig. 9). An AUC calculation was used to represent the total amount of growth over the entire assay. The condition to which no iron source was added, as expected, demonstrated the least amount of growth while the highest amount of growth was detected when hHb was provided. The A+B+ strain was similarly able to utilize hHb, pHb and rHb for growth in this assay. However, mHb supported growth of A+B+ to some extent, with levels significantly higher than the no iron control but growth was significantly lower than detected in the hHb condition. In contrast to the published observations with other gonococcal nutrient transport systems, these results indicate that *N. gonorrhoeae* can bind to and utilize hemoglobin from different species, in addition to human.

**Figure 9:**
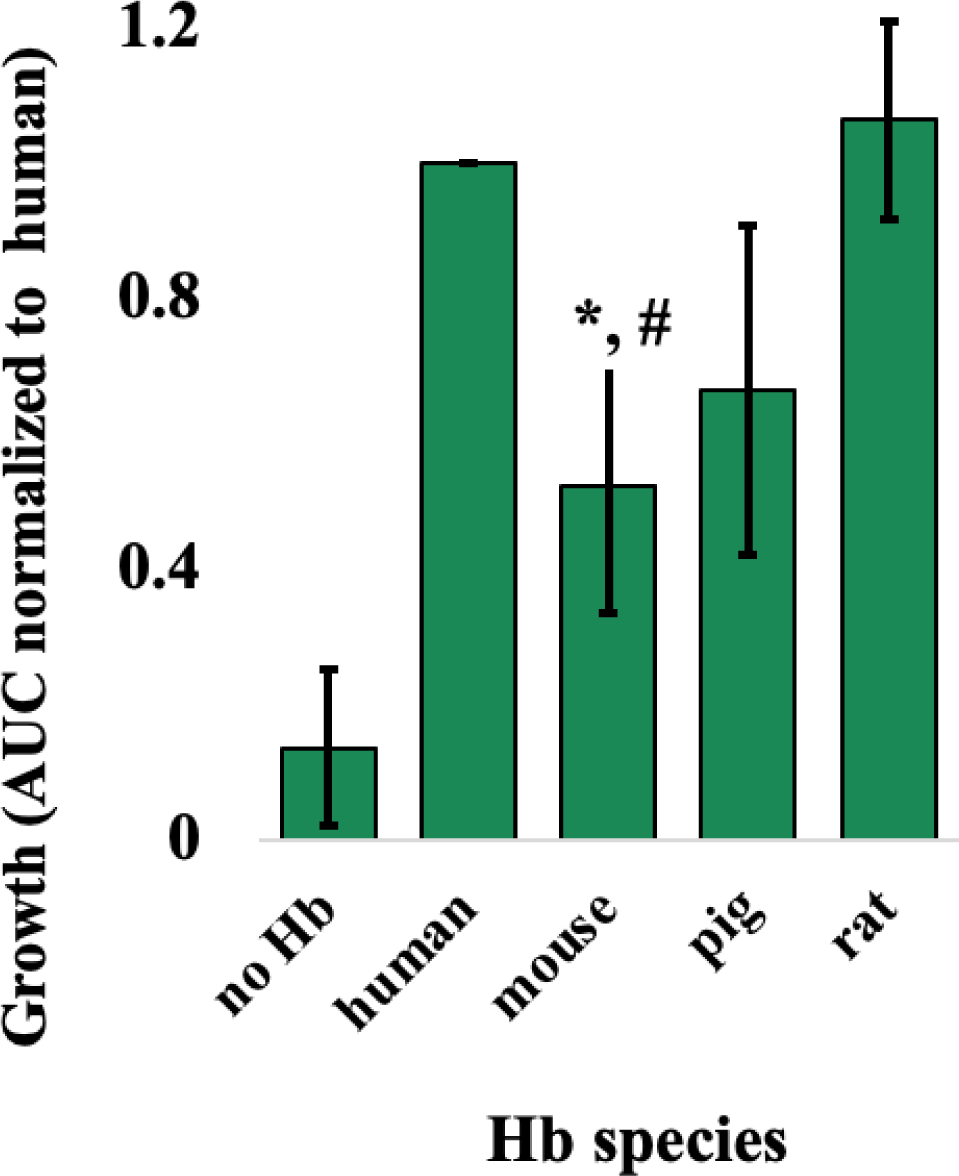
*N. gonorrhoeae* can grow on hemoglobin, produced by species other than human, as a sole iron source. A growth assay was used to establish whether *N. gonorrhoeae* could grow on different Hb species, as their sole iron source. A+B+ was grown on GCB/DFO/ IPTG plates before being resuspended into CDM. The cell suspension was standardized to an OD_600_ of 0.002 before being added to a 96-well plate containing DFO and IPTG with no Hb or 1 µM human, mouse, pig, or rat Hb. Cells were grown for 21 hours while the OD_600_ was recorded in 30 minutes intervals to assess the growth of A+B+ with hemoglobin of different species. The human Hb condition represents the positive control and the no Hb condition represents the negative control. From the growth curves, the area under the curve (AUC) was calculated using GraphPad Prism then normalized to growth on human Hb. Three biological replicates are represented with their means and standard deviation shown. Statistically significant differences were assessed by the Student’s *t-test* relative to the human Hb condition (*, p < 0.05) and no Hb (#, p < 0.05).

## DISCUSSION

The fight against gonorrhea, a serious public health concern, persists. There is no protective immunity elicited following infection; moreover, the number of gonococcal isolates resistant to all available antibiotics is dramatically increasing. The lack of an efficacious vaccine against *N. gonorrhoeae* does not help our odds at beating this pathogen; therefore, there is an urgency to develop an effective vaccine that could significantly prevent and decrease disease incidence. Like other Gram-negative bacteria, *N. gonorrhoeae* requires metal inside the human host but is confronted with nutritional immunity imposed by the host. The human host restricts essential nutrients, like iron, to prevent bacteria from colonizing and causing disease. To overcome these metal-limited conditions imposed by the host, *N. gonorrhoeae* expresses eight outer membrane TdTs that bind to host metal-sequestering proteins to bypass nutritional immunity and access their metal stores (26); of these TdTs, three are bipartite receptors consisting of a surface lipoprotein and a transmembrane protein. The first two are: TbpAB, and the lactoferrin binding protein receptor system (LbpAB) which respectively enable iron uptake from transferrin and lactoferrin (38, 45-48). The third: HpuAB, the subject of the current investigation facilitates iron acquisition from hemoglobin (30, 37).

In our study, we hypothesized that deletion of select extracellular loops from HpuB would reduce or abrogate Hb binding and consequently cause defects in growth with Hb as a sole iron source. We expected that these deletion mutations would help identify domains of the on the HpuB transporter that are essential for ligand binding and iron uptake functions. Loops 2, 3, 4, and 7 of HpuB were targeted in the current study; of note, loop 7 contains conserved motifs found in many heme transporters. These loops were selected due to homology with similar ligand binding domains in TbpA (32, 49-52) and in the meningococcal Hb receptor, HmbR (27, 53). TbpA and HmbR are approximately 25% and 28% identical to HpuB, respectively.

Both HpuA and HpuB were thought to be required for Hb utilization (29, 30, 54). In our study, we generated HpuB loop deletion mutants, near the N-terminus between residues 236 and 559. Surprisingly, we observed that RSC275 (A+B-), expressing only the surface lipoprotein HpuA, could neither bind Hb nor utilize it for growth, whereas RSC276 (A-B+), expressing only transmembrane protein HpuB, could bind hHb but not utilize it as a sole iron source. This suggests a potential role for HpuA in increasing the efficiency of iron internalization by HpuB, as seen with the surface lipoprotein TbpB in the TbpAB system (55).

In the absence of the lipoprotein HpuA, mutant strains with deletions of D236-246 and D306-311 were able to bind to and utilize hHb at A+B+ control levels while D366-370 bound to and utilized Hb at levels significantly lower than that of the A+B+ positive control. This is interesting as RSC276, which also does not express HpuA, cannot utilize hHb when producing the WT version of HpuB, suggesting that these mutations allowed for the need of HpuA to be bypassed during internalization.

In a study by Chen *et al.,* (56) a strain lacking HpuA and producing WT HpuB was unable to grow on hHb (Hb-). However, Hb+ revertants were isolated when this strain was incubated on Hb as a sole source of iron. These revertants resulted from single point mutations in HpuB (clustered towards the C-terminus) and demonstrated restored growth on Hb as a sole iron source, even in the absence of HpuA. Chen *et al.,* attributed this restored growth to the presence of free heme, released from Hb, as addition of human serum albumin prevented the iron uptake phenotype. In their work, they also suggested that the requirement for HpuA could be bypassed by the point mutations in HpuB. The authors suggested that free heme could be imported through the mutated forms of HpuB more readily than the wild-type versions. In the current study, we identified deletion mutations in HpuB that rendered the system competent for heme-iron extraction and import in the absence of HpuA, suggesting that perhaps the function of HpuA is to enable easier access of HpuB to Hb to enhance extraction and subsequent import of heme-iron.

Our study demonstrated that, when HpuA was expressed, D236-246, D306-311 and D366-370 were all able to bind and grow on hHb, with D236-246 growing significantly less and D366-370 binding significantly less than A+B+. Additionally, when looking closely at the AlphaFold2 models, the locations of the loop 2, 3, and 4 deletions all seemed to be near the hHb and/or HpuA interface. Therefore, a possible explanation for the ability of the strains lacking HpuA to grow on hHb could be that the mutations in HpuB circumvent the requirement for HpuA, by making the core of the beta-barrel more accessible and allowing for more efficient heme internalization. We speculate that a potential mechanism to explain our results is that, during hHb binding and metal internalization, HpuA acts on the HpuB extracellular loops by pulling them away from the orifice of the beta-barrel to facilitate more efficient heme uptake, as seen by the enhanced growth phenotypes when HpuA was present. The HpuB deletion mutants might have enough conformational change that shifted the position of these extracellular loops to expose the beta-barrel and enable heme uptake even in the absence of HpuA.

In contrast, the HpuB loop 7 deletion mutants demonstrated abrogated binding and growth on hHb when HpuA was not expressed. Interestingly, when HpuA was expressed, these loop 7 mutants completely recovered their hHb binding abilities; we observed partially recovered growth on hHb although this was still significantly reduced compared to the control A+B+ strain. This points to the importance of loop 7 residues, 538-544 (hypothetical helix), H548 (histidine motif) and 555-559 (NPEL motif), for growth on hHb, but also implicate that HpuA can facilitate the binding of hHb with HpuB in these loop 7 mutants.

Many heme uptake systems in Gram-negative bacteria contain heme-binding motifs, including two conserved histidine residues located between a FRAP and a NXXL motif (42). The histidine residues are not found in all Gram-negatives and examining the sequence from *N. gonorrhoeae* strain FA19, we identified only one histidine residue between a FRAP and a NPEL motif. AlphaFold 2 prediction models of Hb-HpuB with and without HpuA were also generated in this study. The distance between the identified histidine at position 548 (H548) and the heme group of the closest Hb subunit, thought to be the heme source, was measured in the models. In the presence of HpuA, the distance between H548 and the heme group is shorter than when HpuA is absent. This could explain a possible role for HpuA, which is to help stabilize HpuB in a conformation that favors a better binding affinity and shortens the distance between H548 and the heme to be transported across the barrel. This would be different from the other bipartite systems (TbpAB and LbpAB) where the anchored surface lipoprotein directly captures its cognate substrate and facilitates transfer in complex with the barrel protein. Interestingly, regardless of HpuA expression, when comparing distance between H548 and the heme in intact HpuB to that of HpuB D538-544 (hypothetical helix) or HpuB D555-559 (NPEL motif), the distance is longer in the deletion mutants where H548 seem to be pointing away from the surface and into the barrel. This could explain the growth defect seen in the loop 7 mutants as H548 would be too far away to interact with hHb, possibly preventing uptake. Finally, as the presence of HpuA restored the binding capability of the loop 7 mutants, another possible role for HpuA is to aid in ligand binding. One possible mechanism of function could be that HpuA helps orient hHB to stabilize its binding with HpuB, and thereby bringing the heme group close enough to the putative heme binding motif, H548, to initiate heme uptake.

In a previous study, it was noted that although heme did not compete with ^125^I-labelled hHb for binding the meningococcal HpuAB, this system did not discriminate between Hb of different species, possibly due to HpuAB specifically recognizing the heme moiety of Hb (52). A study of meningococcal HmbR also showed a significant difference in the abilities of some animal Hbs to serve as iron sources or to enable the growth stimulation accomplished by hHb (57). In our gonococcal study, the wild-type A+B+ strain was able to bind HRP-conjugated hHb and the binding was also inhibited by unlabeled human, mouse, pig, or rat Hb species, indicating that the gonococcal receptor system, like that described for *N. meningitidis* recognized other species of Hb other than that from humans. Subsequently, we determined that Hb from these different species also allowed growth of the wild-type strain at levels lower than that generated with hHb, but the growth with the alternative forms of Hb still resulted in growth significantly above background. These results are interesting as precedent suggested that *N. gonorrhoeae* exclusively bound to the human forms of the nutritional immunity proteins, including human transferrin, calprotectin and S100A7 (43, 44, 58). However, the sequence conservation of Hbs from the different species is very high, consistent with the ability of the HpuAB system to recognize multiple forms of Hb from distinct animal species. While transgenic mouse models of disease are being developed (59, 60), to assess the contributions of TbpAB and the zinc transport systems to virulence, this may not be necessary for the HpuAB system as the mouse Hb can presumably be employed as an iron and heme source if it is available during experimental infections.

Although most of the gonococci isolated from humans do not express the hemoglobin receptor, the genes encoding the HpuAB system are maintained (30, 37) and could have a role in pathogenesis. One study concluded that the hemoglobin receptors facilitate invasion and dissemination of *N. meningitidis* in the vascular system (61). Moreover, during infection in the early phases of a woman’s menstrual cycle, expression of the gonococcal Hb receptor appears to have a selective advantage. (31). Therefore, elucidating the advantages of expressing these hemoglobin receptors would be crucial in determining their potential as vaccine components.

In summary, this study characterized structure-function relationships in the gonococcal TonB-dependent transporter, HpuB. We showed that some mutations in HpuB bypassed the need for the surface lipoprotein, HpuA, for binding and uptake. We also demonstrated the particular importance of the putative heme motifs, in loop 7, for the internalization of heme. Our findings also highlighted the importance of HpuA, for binding hHb, as the lipoprotein completely rescued the binding defects of the loop 7 mutants. We demonstrated the importance of residues 538-544 which constitute a hypothetical α-helical region in loop 7. This region could have a significant role in the interaction of HpuB with hHb, and in ensuring heme extraction from Hb as seen previously with loop 3 helices of both TbpA and TdfJ (43, 62, 63). Therefore, acquiring a high-resolution structure of HpuB, either by protein x-ray crystallography or cryo-electron microcopy, will help better inform the choice of future mutational targets on the extracellular loops as in other studies (43, 62, 63). As previously published, Transferrin-Binding Protein B (TbpB) and factor H binding protein (fHbp) mutants that are unable to bind host ligands demonstrated better vaccine potential (33-35). Although a similarly enhanced immunogenicity has not yet been described for the TdTs following mutagenesis, it is hypothesized that, a cocktail of non-binding mutants from the TdTs, could enhance vaccine potential and offer protection. Therefore, further characterizing HpuB and other TdTs for vulnerable surface-exposed, ligand-binding motifs is an important step towards potentially developing a protective anti-gonococcal vaccine.

## MATERIALS AND METHODS

### Gonococcal strains growth conditions

Gonococci were propagated on GC medium base (GCB; Difco) agar containing Kellogg’s supplement I (64) and 12.4 µM Fe(NO_3_)_3_ at 36°C with 5% atmospheric CO_2_. In this study, most GCB plates were supplemented with 1 mM Isopropyl-β-D-thiogalactoside (IPTG) and 12.5 µM deferoxamine mesylate (Desferal/DFO) lacking Fe(NO_3_)_3,_ to deprive gonococcal strains of iron. As both HpuA and HpuB are required for internalization, the iron-repressed HpuAB promoter is derepressed by the addition of DFO; and the *lac* promoter described in the construct below, is derepressed by the addition of IPTG. Consequently, the addition of DFO and IPTG allow native expression *hpuA* and ectopic expression of *hpuB*, respectively. The GCB/DFO/IPTG conditions were used to culture GC unless otherwise noted. For growth experiments, cells cultured from GCB/DFO/IPTG plates were inoculated into Chelex (BioRad)-treated defined medium (CDM) at a starting OD_600_ of 0.002. Then the cells were grown, up to 24 hours, with additional DFO, IPTG and hemoglobin (Hb), and optical density readings were taken every hour in a Cytation 5 BioTek microplate reader.

### Gonococcal mutant construction

Table 1 summarizes the different strains and plasmids used and generated in this study. The *hpuB* mutants, with or without native *hpuA*, were generated from wild-type (WT) or mutated *hpuB* gene sequences (from strain FA19) synthesized by Genewiz Inc. using de novo synthesis. The regions mutated were in putative loops 2, 3, 4 and 7, the latter of which contained conserved motifs found in all hemoglobin receptors. These genes were subsequently subcloned into the SmaI site of pVCU234, using restriction endonucleases from New England BioLabs (NEB). These plasmids were then linearized with PciI and used to transform piliated strains RSC150 (*hpuA-hpuB-*) and RSC275 (*hpuA-hpuB*+). GCB plates supplemented with 1 µg/ mL chloramphenicol were used to select for transformants. PCR amplification of lysed chloramphenicol-resistant transformants, followed by sequencing of the resulting amplicons confirmed the sequences of the integrated *hpuB* genes. Sequences of primers and plasmid maps are available upon request.

### HpuB peptide-specific, guinea pig, polyclonal antiserum

The amino acid sequence of HpuB was analyzed to identify potential regions to target for antibody generation. Peptide ^420^NSDYSYFAKLYDPK^433^ was chosen based on its surface accessibility (in extracellular loop 5) and immunogenic profile using the Hopp-Woods algorithm. The HpuB peptide was synthesized using standard Fmoc solid-state peptide chemistry and conjugated to KLH, then emulsified 1:1 (volume) with Freund’s adjuvants for immunization. Two guinea pigs were immunized subcutaneously over a period of ∼10 weeks, with terminal bleeds taken at the end. Peptide and polyclonal antiserum were generated by Biosynth.

### Western blots

Gonococcal suspensions, in phosphate buffer saline (PBS), were normalized before being pelleted, resuspended in 2X Laemmli solubilizing buffer (BioRad), and stored at -20°C. Subsequently, the whole-cell lysates were thawed, mixed with 5% β -mercaptoethanol, and boiled for 5 min. Precast 4 to 20% gradient polyacrylamide gels (Bio-Rad) were used to separate the protein samples, then a stain free image was taken to show equal protein loading in each lane. Proteins were transferred onto a 0.45 µm nitrocellulose membrane (VWR), by electroblot. Next, the blots were blocked in 5% (wt/vol) bovine serum albumin (BSA) dissolved in Tris Buffered Saline with 0.05% Tween 20 (TBST). The blots were then probed with the HpuB peptide-specific guinea pig polyclonal antiserum described above (1:7000 in blocker) for 1 h at room temperature (RT). Next, the blots were washed three times with TBST and then probed with AP-conjugated anti-guinea pig IgG secondary antibody (1:10,000 in blocker) for 1 h. Finally, the blots were washed again using TBST and developed using AP-reactive Nitrobluetetrazolium (NBT) 5-bromo-4-chloro-3-indolylphosphate (BCIP) tablets (Sigma). A final image of the blots was taken on a Bio-Rad ChemiDoc gel imaging system using colorimetric detection.

### Iron-restricted growth with human hemoglobin (hHb)

Adapted from previously described methods (65), strains were streaked onto GCB/DFO/IPTG and incubated at 36°C with 5% CO_2_ for 16-19 hours. To test Hb utilization, iron-starved strains were inoculated at OD_600_ of 0.002 in 1X CDM and were added to a 96-well plate. Wells in the plate already contained: 5 μM DFO, 1 mM IPTG, 5 mM mannitol and 6 mM NaHCO₃ with or without 1 μM hHb. This assay was also used to test the ability of *N. gonorrhoeae* to utilize and grow with Hb of different species, this time the wells in the plate contained 5 μM DFO, 1 mM IPTG, 5 mM mannitol, 6 mM NaHCO₃ and either 1 μM hHb, mHb, pHb or rHb. BioTek Synergy plate reader was used to incubate the cultures grown at 36°C, with shaking with 5% CO_2_ and to measure the OD_600_ every 30 min for up to 24 hours. The area under the curve (AUC) of the resulting growth curves was calculated using GraphPad Prism to compare the growth of the strains. A Student’s *t* test on the AUC was performed for 3 biological replicates in Excel.

### Whole-cell enzyme-linked immunosorbent assay (ELISA)

Gonococci were iron stressed by overnight growth on GC/DFO/IPTG agar plates. Whole gonococcal cells were collected from these plates, resuspended in PBS and standardized to an optical density of 1 (OD_600_). Forty-three microliters of the cell suspension were applied in triplicate for each strain and allowed to dry in a MaxiSorp microtiter dishes (Nunc) plate overnight. The dried microtiter plate was then blocked with 200 µL of 5% (wt/vol) non-fat dry milk in TBS, added for 1 h of incubation. After the blocker was removed, 4 nM HRP-conjugated hHb in blocker was added for 1 h of incubation at RT, after which the cells were then washed five times with TBS. Later, 43 µL of 3,3′,5,5′-Tetramethylbenzidine (TMB) ELISA substrate solution (Thermo) was added to detect the amount of HRP-hHb bound to the cells in each well. After colorization, 43 µL of 0.18 M sulfuric acid (H_2_SO_4_) was added to each well to stop the reaction. Coloration was quantified by reading the absorbance at 450 nm using a Cytation5 plate reader (BioTek). Statistical analysis was performed using Excel. A Student’s *t* test was performed on 3 biological replicates to assess statistical significance.

### Surface exposure testing using trypsin digestion

Protease accessibility assays, following a protocol previously described (38), were done using trypsin. Iron stressed, IPTG treated gonococcal cells were suspended to an optical density OD_600_ of ∼0.4. For 0, 10, 20, 30, and 40 min at 36°C with 5% CO_2_, iron-stressed whole gonococcal cells were treated with 5 μg trypsin (Sigma) per mL of culture. At the appropriate time points, 0.6 trypsin-inhibiting units of aprotinin (Sigma) were used to stop the reaction. The resulting trypsin-digested whole cells were pelleted and resuspended in 2X Laemmli solubilizing buffer. As described above, western blotting was conducted to detect HpuB fragments.

### Protein complex prediction by Alphafold2 and ColabFold

For all protein complex predictions, the Google Colaboratory version of the AlphaFold2 (39) software, ColabFold (41), was used with default settings (model type=alphafold2_multimer_v3, num_recycles=3, num_relax=0). Models were generated using the multimer mode (40) for the ternary complex of HpuAB and the tetrameric hemoglobin, the binary complex of HpuB and hemoglobin, and the different HpuB mutant variants generated in this study. Since ColabFold currently cannot incorporate ligands and cofactors into their structural predictions, the known crystal structure of the oxygenated hemoglobin (PDB:1HHO) was superimposed onto the model to correctly position the heme group within the hemoglobin. Visualization and images were prepared using Pymol (Schrödinger LLC).

### Competition ELISA

RSC284 was iron stressed by overnight growth on DFO plates and resuspended as described above. Whole cells were standardized to an optical density of 0.4 (OD_600_). In the MaxiSorp microtiter dish (Nunc), 100 µL of the cell suspension was applied in quadruplet for each condition to be tested and allowed to dry in plate for 2 days. The dried microtiter plate was also blocked with 200 µL of 5% (wt/vol) non-fat dry milk in TBS. One hundred microliters of each of the following conditions, incubated for 1 hr at RT, was added: blocker only as a negative control, 5 nM HRP-hHb as a positive control and 5 nM HRP-hHb with 20 X excess competitor. Unlabeled human, mouse, porcine and rat Hb were used as competitors. The wells were then washed and developed as described above. This time, the decrease in optical density (due to presence of competitor, unlabeled Hbs) was quantified by reading the absorbance at 450 nm using a Cytation5 plate reader (BioTek). Statistical analysis was performed using Excel and a Student’s *t* test was performed on 4 biological replicates.

## Acknowledgements

We acknowledge Scott Lewis and the immunology department at Biosynth for the generation of the HpuB antiserum. This work was supported by funding from the National Health Service grants R01 A1 AI125421, R01 AI127793, and U19 AI144182. The funders had no role in study design, data collection and analysis, decision to publish or preparation of the manuscript.

## Notes

### Competing Interest Statement

The authors have declared no competing interest.

## REFERENCES

1. WHO. 2021. Gonorrhoea: latest antimicrobial global surveillance results and guidance for vaccine development published. https://www.who.int/news/item/22-11-2021-gonorrhoea-antimicrobial-resistance-results-and-guidance-vaccine-development#:~:text=WHO%20estimates%20that%2082.4%20million,curable%20when%20treated%20with%20antibiotics.

2. CDC. 2023. National Overview of STDs, 2021. Prevention CfDCa, https://www.cdc.gov/std/statistics/2021/overview.htm#Gonorrhea.

3. CDC. 2021. Sexually Transmitted Infections Prevalence, Incidence, and Cost Estimates in the United States. https://www.cdc.gov/std/statistics/prevalence-2020-at-a-glance.htm.

4. Schmidt KA, Schneider H, Lindstrom JA, Boslego JW, Warren RA, Van de Verg L, Deal CD, McClain JB, Griffiss JM. 2001. Experimental gonococcal urethritis and reinfection with homologous gonococci in male volunteers. Sex Transm Dis 28:555–64.

5. Walker CK, Sweet RL. 2011. Gonorrhea infection in women: prevalence, effects, screening, and management. Int J Womens Health 3:197–206.

6. Portnoy J, Mendelson J, Clecner B, Heisler L. 1974. Asymptomatic gonorrhea in the male. Can Med Assoc J 110:169 passim.

7. de Campos FP, Kawabata VS, Bittencourt MS, Lovisolo SM, Felipe-Silva A, de Lemos AP. 2016. Gonococcal endocarditis: an ever-present threat. Autops Case Rep 6:19–25.

8. Anan TJ, Culik DA. 1989. Neisseria gonorrhoeae dissemination and gonococcal meningitis. J Am Board Fam Pract 2:123–5.

9. Quillin SJ, Seifert HS. 2018. Neisseria gonorrhoeae host adaptation and pathogenesis. Nat Rev Microbiol 16:226–240.

10. Liu Y, Feinen B, Russell MW. 2011. New concepts in immunity to Neisseria gonorrhoeae: innate responses and suppression of adaptive immunity favor the pathogen, not the host. Front Microbiol 2:52.

11. Liu Y, Liu W, Russell MW. 2014. Suppression of host adaptive immune responses by Neisseria gonorrhoeae: role of interleukin 10 and type 1 regulatory T cells. Mucosal Immunol 7:165–76.

12. Organization WH. 2021. Gonorrhoea: latest antimicrobial global surveillance results and guidance for vaccine development published. https://www.who.int/news/item/22-11-2021-gonorrhoea-antimicrobial-resistance-results-and-guidance-vaccine-development#:~:text=WHO%20estimates%20that%2082.4%20million,curable%20when%20treated%20with%20antibiotics. Accessed

13. Ohnishi M, Golparian D, Shimuta K, Saika T, Hoshina S, Iwasaku K, Nakayama S, Kitawaki J, Unemo M. 2011. Is Neisseria gonorrhoeae initiating a future era of untreatable gonorrhea?: detailed characterization of the first strain with high-level resistance to ceftriaxone. Antimicrob Agents Chemother 55:3538–45.

14. Unemo M, Shafer WM. 2011. Antibiotic resistance in Neisseria gonorrhoeae: origin, evolution, and lessons learned for the future. Ann N Y Acad Sci 1230:E19–28.

15. Unemo M, Shafer WM. 2014. Antimicrobial resistance in Neisseria gonorrhoeae in the 21st century: past, evolution, and future. Clin Microbiol Rev 27:587–613.

16. St Cyr S, Barbee L, Workowski KA, Bachmann LH, Pham C, Schlanger K, Torrone E, Weinstock H, Kersh EN, Thorpe P. 2020. Update to CDC’s Treatment Guidelines for Gonococcal Infection, 2020. MMWR Morb Mortal Wkly Rep 69:1911-1916.

17. Cámara J, Serra J, Ayats J, Bastida T, Carnicer-Pont D, Andreu A, Ardanuy C. 2012. Molecular characterization of two high-level ceftriaxone-resistant Neisseria gonorrhoeae isolates detected in Catalonia, Spain. J Antimicrob Chemother 67:1858–60.

18. Unemo M, Golparian D, Nicholas R, Ohnishi M, Gallay A, Sednaoui P. 2012. High-level cefixime- and ceftriaxone-resistant Neisseria gonorrhoeae in France: novel penA mosaic allele in a successful international clone causes treatment failure. Antimicrob Agents Chemother 56:1273–80.

19. Singh A, Turner JM, Tomberg J, Fedarovich A, Unemo M, Nicholas RA, Davies C. 2020. Mutations in penicillin-binding protein 2 from cephalosporin-resistant Neisseria gonorrhoeae hinder ceftriaxone acylation by restricting protein dynamics. J Biol Chem 295:7529–7543.

20. Fifer H, Livermore DM, Uthayakumaran T, Woodford N, Cole MJ. 2021. What’s left in the cupboard? Older antimicrobials for treating gonorrhoea. J Antimicrob Chemother 76:1215–1220.

21. Russell MW, Jerse AE, Gray-Owen SD. 2019. Progress Toward a Gonococcal Vaccine: The Way Forward. Front Immunol 10:2417.

22. Cornelissen CN, Hollander A. 2011. TonB-Dependent Transporters Expressed by *Neisseria gonorrhoeae*. Front Microbiol 2:117.

23. Cornelissen CN, Anderson JE, Boulton IC, Sparling PF. 2000. Antigenic and sequence diversity in gonococcal transferrin-binding protein A. Infect Immun 68:4725–35.

24. Weinberg ED. 1975. Nutritional immunity. Host’s attempt to withold iron from microbial invaders. Jama 231:39-41.

25. Kehl-Fie TE, Skaar EP. 2010. Nutritional immunity beyond iron: a role for manganese and zinc. Curr Opin Chem Biol 14:218–24.

26. Neumann W, Hadley RC, Nolan EM. 2017. Transition metals at the host-pathogen interface: how Neisseria exploit human metalloproteins for acquiring iron and zinc. Essays Biochem 61:211–223.

27. Lewis LA, Gray E, Wang YP, Roe BA, Dyer DW. 1997. Molecular characterization of hpuAB, the haemoglobin-haptoglobin-utilization operon of Neisseria meningitidis. Mol Microbiol 23:737–49.

28. Lee BC. 1992. Isolation of haemin-binding proteins of Neisseria gonorrhoeae. J Med Microbiol 36:121–7.

29. Lewis LA, Gipson M, Hartman K, Ownbey T, Vaughn J, Dyer DW. 1999. Phase variation of HpuAB and HmbR, two distinct haemoglobin receptors of *Neisseria meningitidis* DNM2. Mol Microbiol 32:977–89.

30. Chen CJ, Elkins C, Sparling PF. 1998. Phase variation of hemoglobin utilization in *Neisseria gonorrhoeae*. Infect Immun 66:987–93.

31. Anderson JE, Leone PA, Miller WC, Chen C, Hobbs MM, Sparling PF. 2001. Selection for expression of the gonococcal hemoglobin receptor during menses. J Infect Dis 184:1621–3.

32. Noto JM, Cornelissen CN. 2008. Identification of TbpA residues required for transferrin-iron utilization by *Neisseria gonorrhoeae*. Infect Immun 76:1960–9.

33. Frandoloso R, Martínez-Martínez S, Calmettes C, Fegan J, Costa E, Curran D, Yu RH, Gutiérrez-Martín CB, Rodríguez-Ferri EF, Moraes TF, Schryvers AB. 2015. Nonbinding site-directed mutants of transferrin binding protein B exhibit enhanced immunogenicity and protective capabilities. Infect Immun 83:1030–8.

34. Martínez-Martínez S, Frandoloso R, Rodríguez-Ferri EF, García-Iglesias MJ, Pérez-Martínez C, Álvarez-Estrada Á, Gutiérrez-Martín CB. 2016. A vaccine based on a mutant transferrin binding protein B of Haemophilus parasuis induces a strong T-helper 2 response and bacterial clearance after experimental infection. Vet Immunol Immunopathol 179:18–25.

35. Beernink PT, Shaughnessy J, Braga EM, Liu Q, Rice PA, Ram S, Granoff DM. 2011. A meningococcal factor H binding protein mutant that eliminates factor H binding enhances protective antibody responses to vaccination. J Immunol 186:3606–14.

36. Liu X, Olczak T, Guo HC, Dixon DW, Genco CA. 2006. Identification of amino acid residues involved in heme binding and hemoprotein utilization in the Porphyromonas gingivalis heme receptor HmuR. Infect Immun 74:1222–32.

37. Chen CJ, Sparling PF, Lewis LA, Dyer DW, Elkins C. 1996. Identification and purification of a hemoglobin-binding outer membrane protein from *Neisseria gonorrhoeae*. Infect Immun 64:5008–14.

38. Cornelissen CN, Sparling PF. 1996. Binding and surface exposure characteristics of the gonococcal transferrin receptor are dependent on both transferrin-binding proteins. J Bacteriol 178:1437–44.

39. Jumper J, Evans R, Pritzel A, Green T, Figurnov M, Ronneberger O, Tunyasuvunakool K, Bates R, Žídek A, Potapenko A, Bridgland A, Meyer C, Kohl SAA, Ballard AJ, Cowie A, Romera-Paredes B, Nikolov S, Jain R, Adler J, Back T, Petersen S, Reiman D, Clancy E, Zielinski M, Steinegger M, Pacholska M, Berghammer T, Bodenstein S, Silver D, Vinyals O, Senior AW, Kavukcuoglu K, Kohli P, Hassabis D. 2021. Highly accurate protein structure prediction with AlphaFold. Nature 596:583–589.

40. Richard E, Michael ON, Alexander P, Natasha A, Andrew S, Tim G, Augustin Ž, Russ B, Sam B, Jason Y, Olaf R, Sebastian B, Michal Z, Alex B, Anna P, Andrew C, Kathryn T, Rishub J, Ellen C, Pushmeet K, John J, Demis H. 2022. Protein complex prediction with AlphaFold-Multimer. bioRxiv:2021.10.04.463034.

41. Mirdita M, Schütze K, Moriwaki Y, Heo L, Ovchinnikov S, Steinegger M. 2022. ColabFold: making protein folding accessible to all. Nat Methods 19:679–682.

42. Bracken CS, Baer MT, Abdur-Rashid A, Helms W, Stojiljkovic I. 1999. Use of heme-protein complexes by the Yersinia enterocolitica HemR receptor: histidine residues are essential for receptor function. J Bacteriol 181:6063–72.

43. Cash DR, Noinaj N, Buchanan SK, Cornelissen CN. 2015. Beyond the Crystal Structure: Insight into the Function and Vaccine Potential of TbpA Expressed by *Neisseria gonorrhoeae*. Infect Immun 83:4438–49.

44. Kammerman MT, Bera A, Wu R, Harrison SA, Maxwell CN, Lundquist K, Noinaj N, Chazin WJ, Cornelissen CN. 2020. Molecular Insight into TdfH-Mediated Zinc Piracy from Human Calprotectin by Neisseria gonorrhoeae. mBio 11.

45. Cornelissen CN, Sparling PF. 1994. Iron piracy: acquisition of transferrin-bound iron by bacterial pathogens. Mol Microbiol 14:843–50.

46. Biswas GD, Sparling PF. 1995. Characterization of lbpA, the structural gene for a lactoferrin receptor in Neisseria gonorrhoeae. Infect Immun 63:2958–67.

47. Biswas GD, Anderson JE, Chen CJ, Cornelissen CN, Sparling PF. 1999. Identification and functional characterization of the Neisseria gonorrhoeae lbpB gene product. Infect Immun 67:455–9.

48. Pettersson A, Prinz T, Umar A, van der Biezen J, Tommassen J. 1998. Molecular characterization of LbpB, the second lactoferrin-binding protein of Neisseria meningitidis. Mol Microbiol 27:599–610.

49. Boulton IC, Yost MK, Anderson JE, Cornelissen CN. 2000. Identification of discrete domains within gonococcal transferrin-binding protein A that are necessary for ligand binding and iron uptake functions. Infect Immun 68:6988–96.

50. Masri HP, Cornelissen CN. 2002. Specific ligand binding attributable to individual epitopes of gonococcal transferrin binding protein A. Infect Immun 70:732–40.

51. Yost-Daljev MK, Cornelissen CN. 2004. Determination of surface-exposed, functional domains of gonococcal transferrin-binding protein A. Infect Immun 72:1775–85.

52. Rohde KH, Gillaspy AF, Hatfield MD, Lewis LA, Dyer DW. 2002. Interactions of haemoglobin with the *Neisseria meningitidis* receptor HpuAB: the role of TonB and an intact proton motive force. Mol Microbiol 43:335–54.

53. Perkins-Balding D, Baer MT, Stojiljkovic I. 2003. Identification of functionally important regions of a haemoglobin receptor from Neisseria meningitidis. Microbiology (Reading) 149:3423–3435.

54. Biswas GD, Anderson JE, Sparling PF. 1997. Cloning and functional characterization of Neisseria gonorrhoeae tonB, exbB and exbD genes. Mol Microbiol 24:169–79.

55. Anderson JE, Sparling PF, Cornelissen CN. 1994. Gonococcal transferrin-binding protein 2 facilitates but is not essential for transferrin utilization. J Bacteriol 176:3162–70.

56. Chen CJ, McLean D, Thomas CE, Anderson JE, Sparling PF. 2002. Point mutations in HpuB enable gonococcal HpuA deletion mutants to grow on hemoglobin. J Bacteriol 184:420–6.

57. Stojiljkovic I, Larson J, Hwa V, Anic S, So M. 1996. HmbR outer membrane receptors of pathogenic Neisseria spp.: iron-regulated, hemoglobin-binding proteins with a high level of primary structure conservation. J Bacteriol 178:4670–8.

58. Maurakis S, Keller K, Maxwell CN, Pereira K, Chazin WJ, Criss AK, Cornelissen CN. 2019. The novel interaction between Neisseria gonorrhoeae TdfJ and human S100A7 allows gonococci to subvert host zinc restriction. PLoS Pathog 15:e1007937.

59. Zarantonelli ML, Szatanik M, Giorgini D, Hong E, Huerre M, Guillou F, Alonso JM, Taha MK. 2007. Transgenic mice expressing human transferrin as a model for meningococcal infection. Infect Immun 75:5609–14.

60. Levy M, Aouiti Trabelsi M, Taha MK. 2020. Evidence for Multi-Organ Infection During Experimental Meningococcal Sepsis due to ST-11 Isolates in Human Transferrin-Transgenic Mice. Microorganisms 8.

61. Harrison OB, Bennett JS, Derrick JP, Maiden MCJ, Bayliss CD. 2013. Distribution and diversity of the haemoglobin-haptoglobin iron-acquisition systems in pathogenic and non-pathogenic Neisseria. Microbiology (Reading) 159:1920–1930.

62. Maurakis SA, Stoudenmire JL, Rymer JK, Chazin WJ, Cornelissen CN. 2022. Mutagenesis of the Loop 3 alpha-Helix of Neisseria gonorrhoeae TdfJ Inhibits S100A7 Binding and Utilization. mBio 13:e0167022.

63. Greenawalt AN, Stoudenmire J, Lundquist K, Noinaj N, Gumbart JC, Cornelissen CN. 2022. Point Mutations in TbpA Abrogate Human Transferrin Binding in Neisseria gonorrhoeae. Infect Immun 90:e0041422.

64. Kellogg DS, Jr., Peacock WL, Jr., Deacon WE, Brown L, Pirkle DI. 1963. Neisseria gonorrhoeae. I. Virulence Genetically Linked to Clonal Variation. J Bacteriol 85:1274–9.

65. Maurakis S, Cornelissen CN. 2020. Metal-Limited Growth of *Neisseria gonorrhoeae* for Characterization of Metal-Responsive Genes and Metal Acquisition from Host Ligands. J Vis Exp.

66. Cash DR. 2016. Drug and Vaccine Development for *Neisseria gonorrhoeaea*. PhD dissertation. Virginia Commonwealth University, Richmond, VA.

