## Supplemental Figures for "Investigating the importance of surface exposed loops in the gonococcal HpuB transporter for hemoglobin binding and utilization"

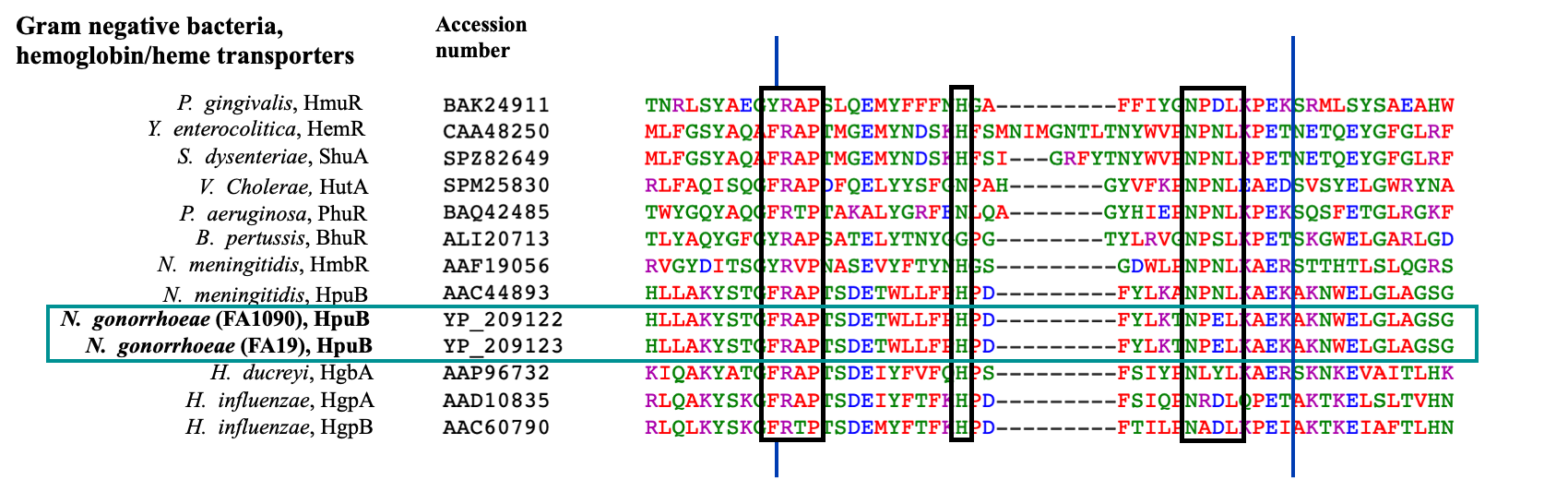

**Supplemental figure 1: Muti-sequence alignment of Gram-negative hemoglobin/heme transporters.** Only loop 7 is shown and it is delineated by the blue lines. The different bacteria used are shown with their corresponding accession number. The conserved FRAP (YRAP and FRTP variation seen), Histidine (sometimes absent) and NXXL motifs are shown in the black rectangles. The amino acid sequence of *N. gonorrhoeae*, with the FRAP, histidine, and NPEL motifs, is shown in the cyan rectangle.

**A	B**

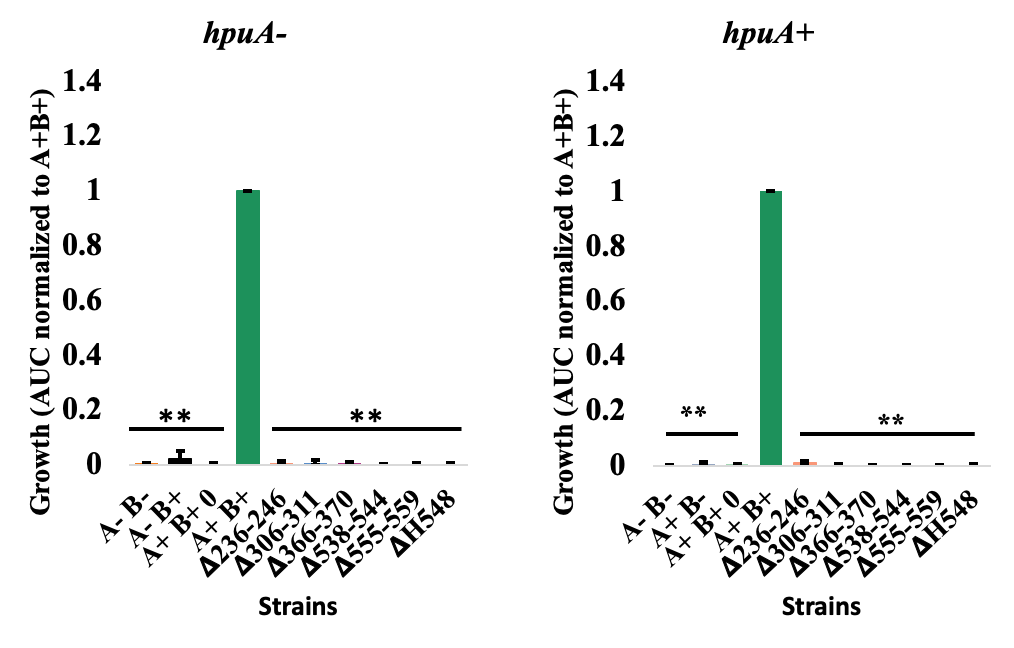

**Supplemental figure 2: HpuB mutants are impaired for growth in the absence of hHb.** Gonococci were grown on GCB/DFO/ IPTG plates before being resuspended into CDM. The cell suspensions were standardized to an OD_600_ of 0.002 before being added to a 96-well plate containing DFO, IPTG and 0 µM hHb. Cells were grown for 21 hours while the OD_600_ was recorded in 30 minutes intervals to assess the growth of strains without native hpuA (A) and strains with native hpuA (B). A-B- is used as a negative control and A+B+ as a positive control with 1 µM Hb. A+B+ 0 µM Hb, A+B- as well as A-B+ also represent controls. The area under the curve (AUC) was calculated using GraphPad Prism then normalized to A+B+. Three biological replicates are represented with their means and standard deviation shown. A Student’s *t-test* was used to calculate the statistical significance relative to A+B+ (**, p < 0.005).

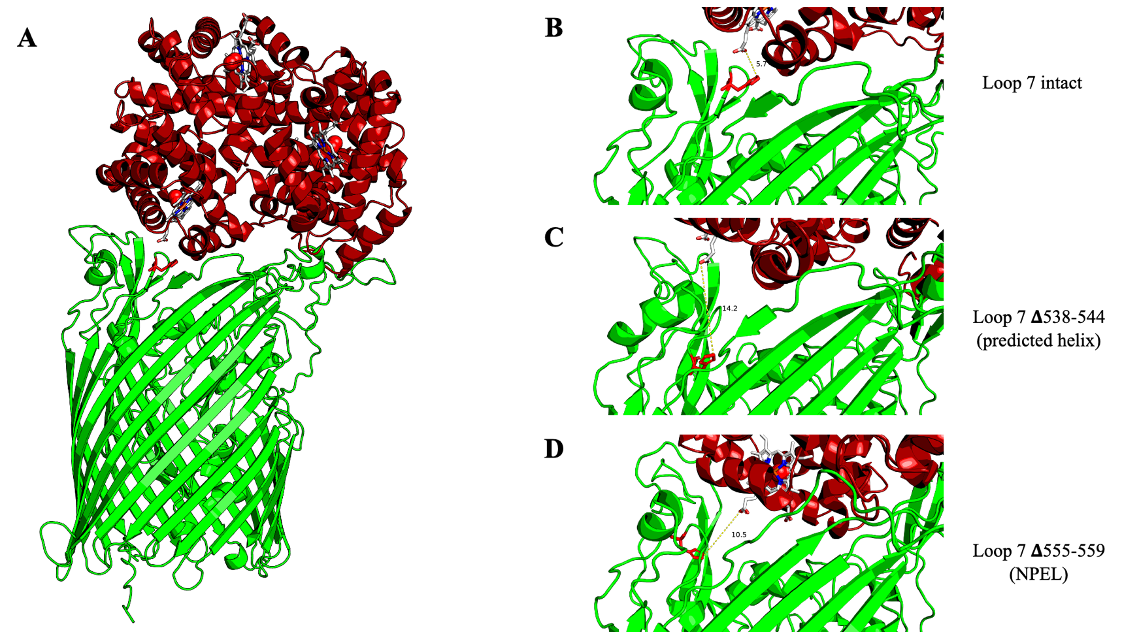

**Supplemental figure 3: The deletions in loop 7 move histidine 548 further away from the closest heme group.** HpuB-Hb structure. AlphaFold2 was used to get a prediction model of the structures. A known Hb-heme crystal structure (PDB:1hho) was super imposed to show the heme groups. HpuB is in green with H548 highlighted in red, Hb in red, heme group in white with is Fe2+ as a red dot. The distance in angstroms (Å) between the heme group and histidine at position 548 is measured.
